# *Nanoviricides* Platform Technology based NV-387 polymer Protects Remdesivir from Plasma-Mediated Catabolism *in vitro:* Importance of its increased lifetime for *in vivo* action

**DOI:** 10.1101/2021.10.22.465399

**Authors:** Ashok Chakraborty, Anil Diwan, Vinod Arora, Yogesh Thakur, Preetam Holkar, Vijetha Chinige

## Abstract

As of today seven coronaviruses were identified to infect humans, out of which only 4 of them belongs to beta family of coronavirus, like HCoV-HKU1, SARS-CoV-2, MERS-CoV and SARS-CoV. SARS family of viruses were known to cause severe respiratory disease in humans. SARS-CoV-2 infection causes pandemic COVID-19 disease with high morbidity and mortality. Remdesivir (RDV) is the only antiviral drug so far approved for Covid-19 therapy by FDA. However it’s efficacy is limited *in vivo* due to it’s low stability in presence of Plasma.

Here we show the stability of RDV encapsulated with our platform technology based polymer NV-387 (NV-CoV-2-R), in presence of Plasma *in vitro* in comparison to naked RDV when incubated in plasma. The potential use of this polymer *in vivo* will be discussed, here.

## Introduction

A new threat to human life, in recent days, is the outbreak of SARS-CoV-2 human corona virus in China, at Wuhan City in late December 2019 **[1]** [Andersen et. al. 2020]. SARS-COV-2 is a beta family of human coronavirus and **stands for “*S****evere Acute Respiratory Syndrome Coronavirus 2”* that causes the severe lower tract infection called COVID-19 **[For rev. 2–4] [**Singhal 2020; Chakraborty et. al. 2020, 2020a]. As of Sept 30, 2021, the world-wide SARS-CoV-2 infected cases are more than 219M and death number is about 4.55M **[5]** [Covid.cdc.gov/covid-data-tracker/; Updated: Sept 12, 2021]. In USA, there have been over 44.5 million confirmed cases with 715K deaths (approx.). **[6]** [<u>WHO Coronavirus Disease (COVID-19) Dashboard</u>. Data last updated: 2021/10/10. https://covid19.who.int/].

The infection capabilities of SARS-CoV-2 are believed to be primarily due to its increased affinity for the angiotensin-converting enzyme 2 (ACE2) receptor on the body’s cell surface **[1, 2] [**Andersen et. al. 2020; Singhal 2020]. We have recently written an article on referring the zoonosis, susceptibility and different strategies to develop therapeutics **[3] [**Chakraborty et. al. 2020].

COVID-19 was declared as a global pandemic on 11 March 2020, by the WHO, however, the evidence for therapies against this virus is as yet inadequate. Various medical teams are prescribing drugs for this collection of ailments based on some mechanistic data but with limited clinical findings in support of their activity. The clinical importance of pre-clinically validated regimens relies on the available pharmacokinetic (PK) data for the particular drug **[1, 3]** [Andersen et. al. 2020; Chakraborty et. al. 2020].

SARS-CoV-2 virus enters a host cells by binding to and fusing with cell membrane receptor, ACE-2, followed by membrane fusion. Once inside, the virus uses the host cell’s machinery to replicate, using the virus’s RNA dependent RNA polymerase (RdRp) for making genome and transcript copies. Among the different strains of the corona virus, this non-structural protein is unique in structure, and thus making it a potentially useful drug target. *Sofosbuvir*, a synthetic analogue of nucleosides and nucleotides which inhibits RdRp has led to a successful treatment for hepatitis C infection **[7] [**Xie et. al. 2016]. Based on these facts, *remdesivir* was approved by FDA in various clinical trials for the treatment of COVID-19 infected people **[8]** [COVID-19 Treatment Guidelines. 2020. https://www.covid19treatmentguidelines.nih.gov/antiviral-therapy/remdesivir/].

*Remdesivir*, formerly known as GS-5734, is a nucleotide analogue that is claimed to have been originally developed as a treatment against Ebola **[3]**. This drug can also inhibit corona virus replication by inhibiting RNA polymerases (RdRp4). This compound has shown broad antiviral activity *in vitro* against Middle East respiratory syndrome coronavirus (MERS-CoV), severe acute respiratory syndrome coronavirus 1 (SARS-CoV-1), and SARS-CoV-2 **[9–11] [**Gordon et. al. 2020; Sheahan et. al. 2017; Wang et. al. 2020].

In animal studies, *remdesivir* has been found effective in protecting rhesus monkeys from MERS-CoV infection, when given prior to infection **[12]** de Wit E, et. al. 2020]. It also protected African green monkeys from Nipah virus, a cause of fatal encephalitis; and also rhesus monkeys from Ebola virus **[13, 14] [**Lo MK, et. al. 2019; Warren TK, et. al. 2016]. A randomized, well marked, controlled animal study with 12 rhesus monkeys infected with SARS-CoV-2 reported that an attenuation of respiratory symptoms and reduction in lung damage with *remdesivir* administered 12 hours after virus infection **[15] [**Williamson. et. al. 2020].

However, there are a lot of limitation of using *remdesivir in* vivo against COVID-19. (i) Efficacy of *remdesivir in vitro* does not match with the clinical outcomes in humans. (ii) Further, *remdesivir* has some side effects. In the Ebola trial, the side effects of *remdesivir* (RDV) were possible liver damage **[16] [**Remdesivir (RDV). 2020. https://www.rxlist.com/consumer_remdesivir_rdv/drugs-condition.htm]; and also affects kidney and mitochondria **[17, 18] [**Adamsick ML, et. al. 2020; Saleh J, et. al. 2020].

Besides, *remdesivir* is not stable in presence of Plasma. Metabolism of RDV decreases the effective concentration of the drug, *in vivo*, and the time of exposure with the drug that may be needed to eliminate the virus.

We searched for the methodologies to protect *remdesivir* from plasma-mediated metabolism. NanoViricides, Inc. (Shelton, CT, USA) is probably the only company that has developed a PEG-based polymer comprised of a polyethylene glycol (PEG-1000) and C16-alkyl pendants in the monomer unit. PEG forms the hydrophilic shell; while the alkyl chains float together to make a flexible core, like an immobilized oil droplet. This polymer can (a) directly attack the virus and disable them from infecting human cells, and simultaneously, and (b) block the reproduction of the virus that has already gone inside a cell **(Fig. 1).** Together, this double-whammy would result in a cure **[3]** [Chakraborty and Diwan, 2020].

**Fig. 1:**
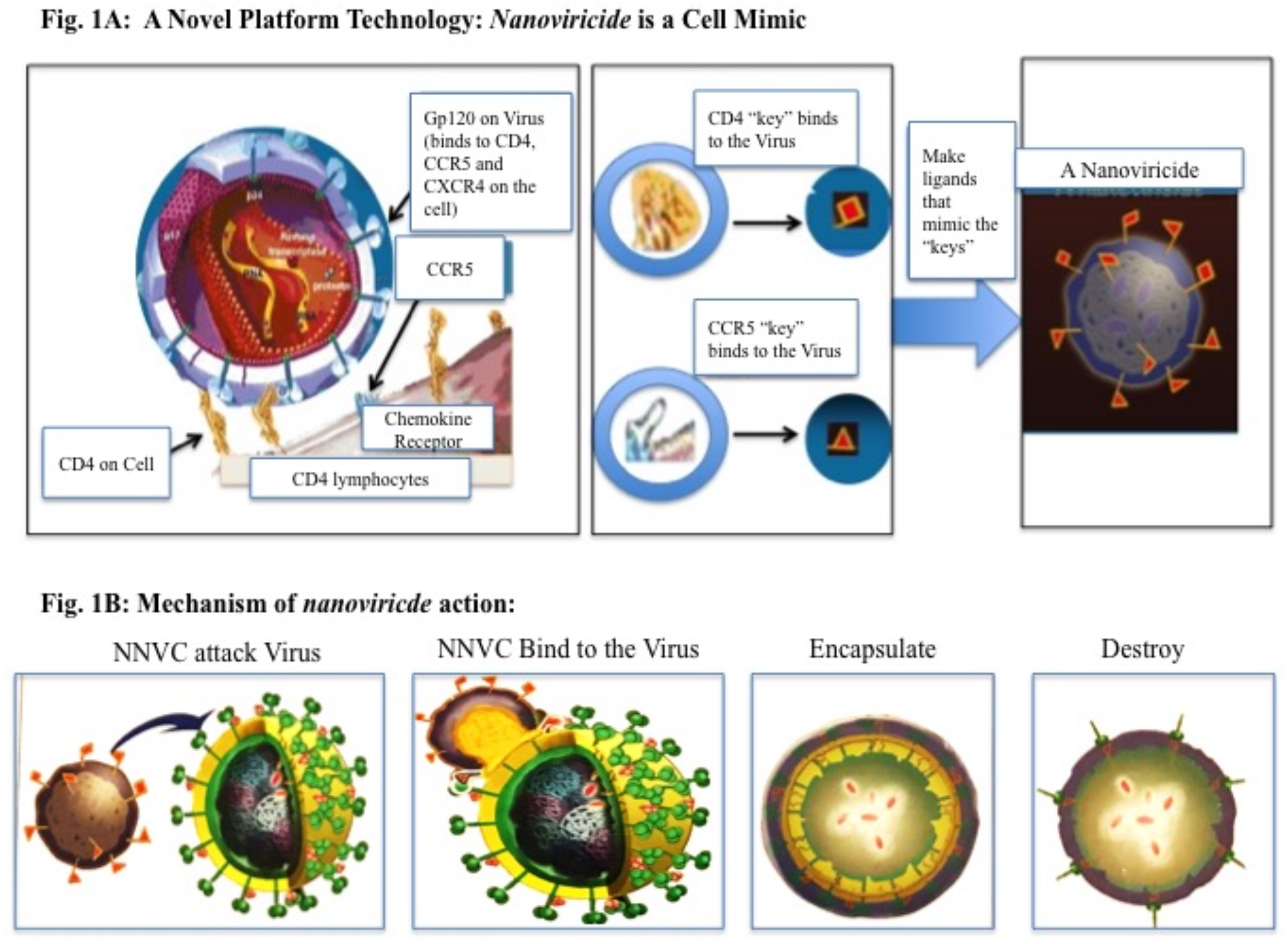
**A: Novel Platform technology: Nanoviricide is a Cell Mimic: A Virus particle binding to a Cell via CD4 and a co-Receptor.** A *nanoviricide* “looks like” a human cell to the virus. *Nanoviricide* is large enough for a virus particle to latch onto it, however, it is yet small enough to circulate readily in the body. Rather than a virus particle entering into a *nanoviricide*, a *nanoviricide* wraps around the virus particle and encapsulate it, by using the virus particle’s very same ability to enter a cell. Viral resistance to the *Nanoviricide* drug is unlikely because even as the virus mutates, it still binds to the same cell surface receptor(s), in the same fashion. **B: A schematic presentation of *nanoviricide* action on virus particle.**

In this communication our objective is to determine the efficacy of the polymer whether can protect RDV from plasma-mediated catabolism. Time dependent RDV level were measured by LC-MS analysis after incubation of the naked RDV vs polymer encapsulated RDV with rat plasma *in vitro*. RDV-metabolite, GS-441524 was also measured to justify the RDV breakdown. For comparison we used commercially available Gilead Remdesivir, and as a negative control we used vehicle (DMSO:MeOH, 1:9). Here we report our results.

## Materials and Methods

### 1.1: Test Articles

**(Table-1.1):**
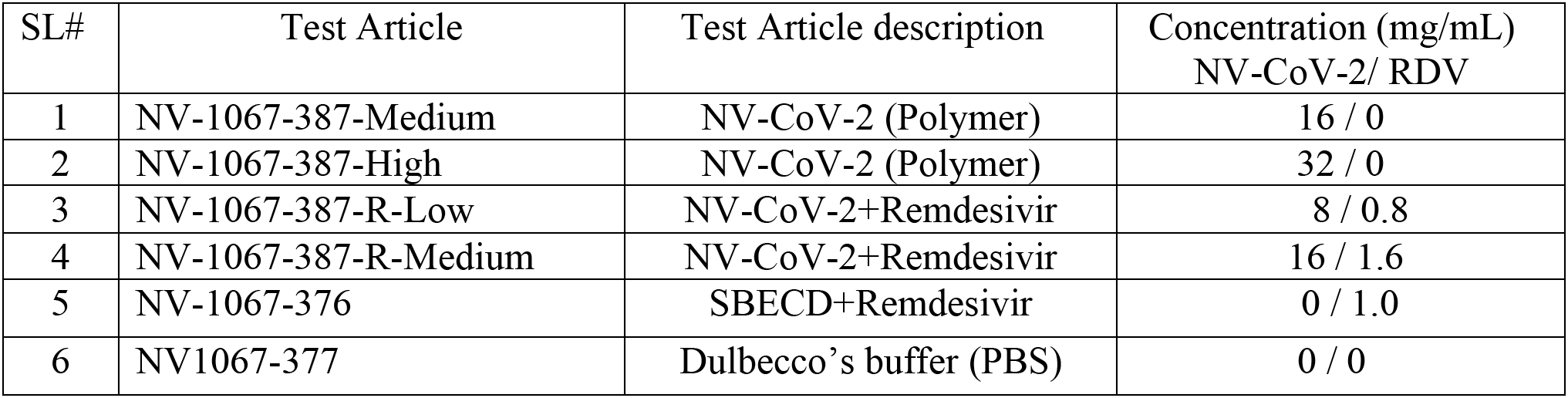

### 1.2: Sources of Reagents

**(Table-1.2):**
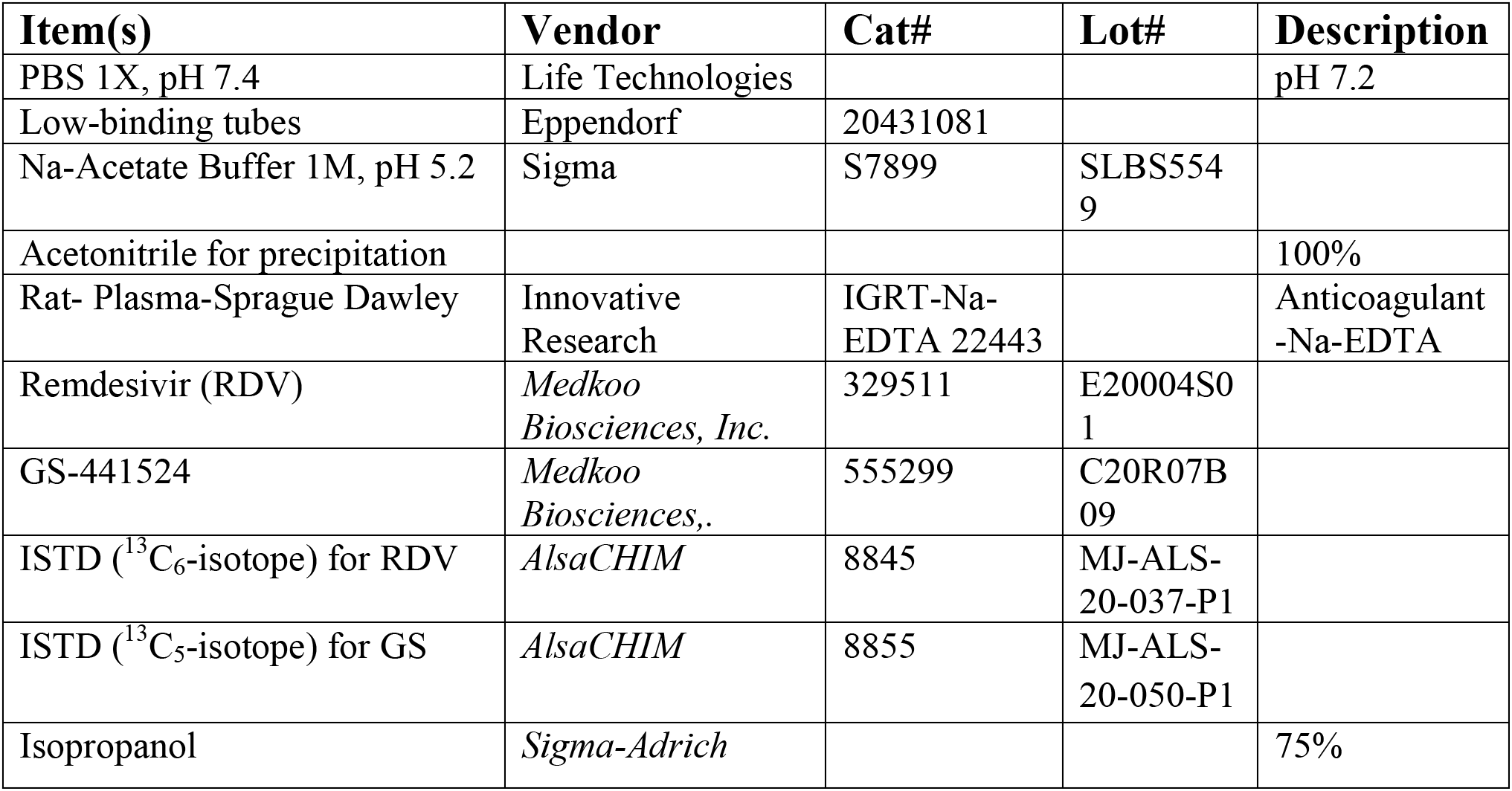

1.2: Sources of Reagents (Table-1.2):
| Item(s) | Vendor | Cat# | Lot# | Description |
| --- | --- | --- | --- | --- |
| PBS 1X, pH 7.4 | Life Technologies |  |  | pH 7.2 |
| Low-binding tubes | Eppendorf | 20431081 |  |  |
| Na-Acetate Buffer 1M, pH 5.2 | Sigma | S7899 | SLBS5549 |  |
| Acetonitrile for precipitation |  |  |  | 100% |
| Rat- Plasma-Sprague Dawley | Innovative Research | IGRT-Na-EDTA 22443 |  | Anticoagulant -Na-EDTA |
| Remdesivir (RDV) | Medkoo Biosciences, Inc. | 329511 | E20004S01 |  |
| GS-441524 | Medkoo Biosciences,. | 555299 | C20R07B09 |  |
| ISTD (<sup>13</sup>C<sub>6</sub>-isotope) for RDV | AlsaCHIM | 8845 | MJ-ALS-20-037-P1 |  |
| ISTD (<sup>13</sup>C<sub>5</sub>-isotope) for GS | AlsaCHIM | 8855 | MJ-ALS-20-050-P1 |  |
| Isopropanol | Sigma-Adrich |  |  | 75% |

### 1.3: Standard Curve Preparation of Remdesivir and GS-441524

were determined by using LC-MS, using different concentration of the standards solution in DMSO + MeOH (1:9) ranging from 0-5 ng/uL final concentrations. Plasma and/or PBS were used in the reaction mixture, as when needed. As an internal standard (ISTD) ^13^C_6_–RDV and ^13^C_5_-GS441524, 2.5 ng/uL each, were used, respectively for RDV and GS assay, and or in mixture. The Final concentration of the internal standards in the reaction mixture becomes 0.125 ng/uL. Extraction of RDV and it’s metabolite GS-441524, for LC-MS assay were done by using Acetonitrile.

Concentrations of RDV and GS-441524 vs. their ratio with the internal standards, ISTDs (^13^C_6_-RDV, ^13^C_5_-GS, respectively) were used to generate a standard curve for RDV and GS-441524, respectively, as shown in the **Table-2 & Figure-2.**

**Fig. 2:**
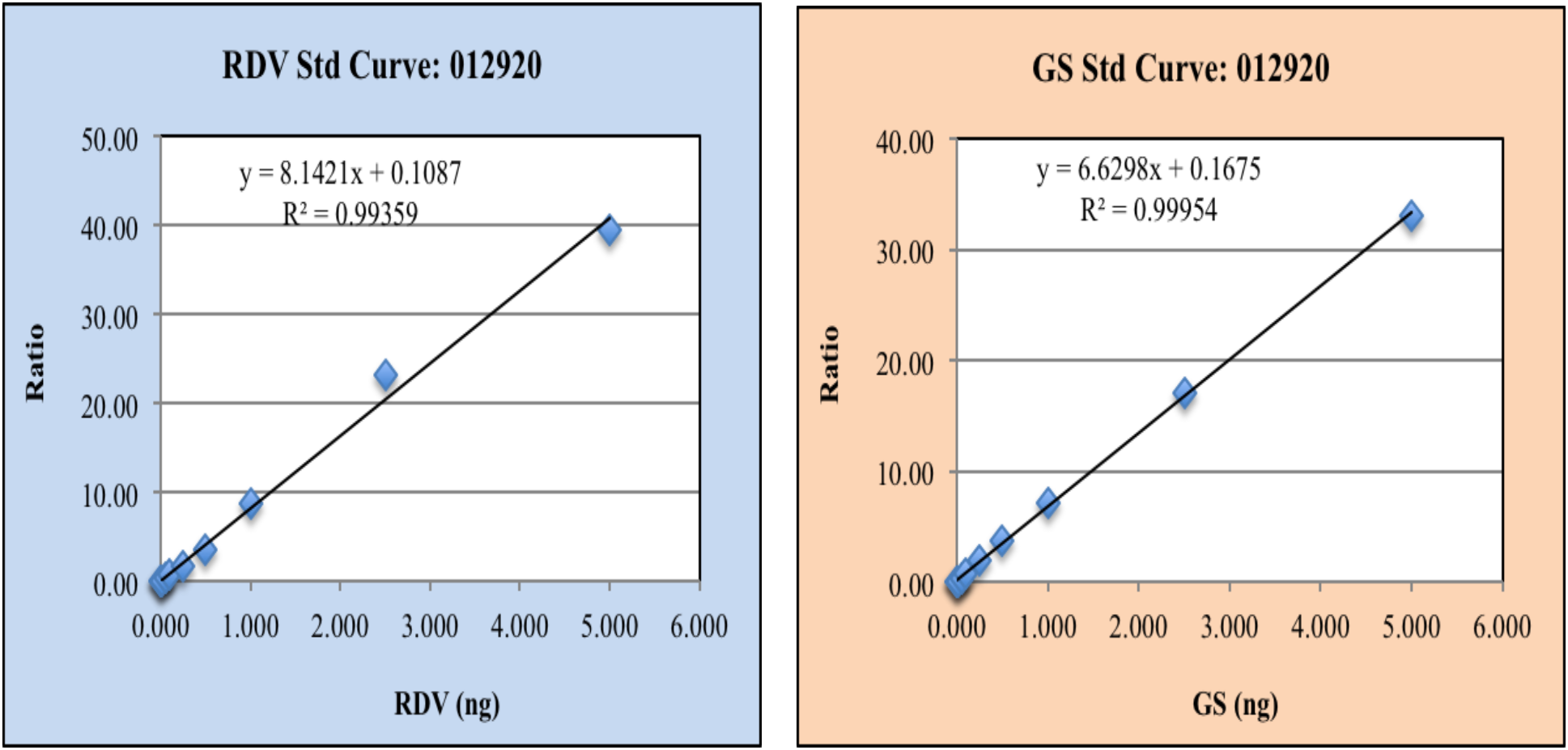
**Standard Curve of Remdesivir and GS-441524** were determined by using LC-MS, using different concentration of the standards solution in DMSO + MeOH (1:9) ranging from 0-5 ug/mL final concentrations. Plasma and/or PBS were used in the reaction mixture, as when needed. As an internal standard (ISTD) ^13^C_6_ – RDV and ^13^C_5_-GS, 2.5 ug/mL each, were used, respectively for RDV and GS assay, and or in mixture. Final extraction of RDV and it’s metabolite GS, were dome by using acetonitrile.

### 1.4: Incubation of test materials with rat plasma (RPL) *in vitro*, and isolation of the RDV/GS-441524 by Acetonitrile

Ten microliter of the test materials (20 ng/uL) were incubated with 30 uL rat plasma for different time points (as indicated in the Figs.), and then extracted with Acetonitrile (100 uL). The mixture is centrifuged for 10 mins at 10,000 × g to separate the supernatants from the precipitates. The supernatants containing either RDV and/or its metabolites, GS-441524 were determined by LC-MS chromatography (as described below).

### 1.5: Detection of RDV and GS-441524 by LC-MS spectroscopy

Analysis was performed using the LC and MS analysis conditions shown in **Table-1.3,** and the multiple reaction monitoring (MRM) data acquisition parameters shown in **Table-1.4**. Shim-Pack Sceptor^™^ C18-120 (50 mm × 22.1 mm I.D., 1.9 uM) was used as the analytical column.

**Table-1.3:** Chromatographic Conditions:

| LC Analysis Conditions |  |
| --- | --- |
| **HPLC Method:** | RDVmetIS_LC_NoSlit-0.4mL-ExtRinse_202010231219.1cm |
| **Column:** | Shimadzu, Shim Pack, Sceptor, C18-120, 2.1x50 mm, 1.9 um, Cat#227-31120-01 |
| **Pre-Column:** | Guard Column, Shimadzu, Shim-pack, C18-120 EXP (G), 1.9 um, 2.1 x 5 mm, or Restek prefilter: Ultrafield UHPLC, 0.2 um frit, Catalog# 25810. |
| **Mobile Phase:** | A: Water containing 0.05% formic acid<br>B: Acetonitrile containing 0.05% formic acid |
| **Gradient:** | 0-0.3 min: 5% B,<br>0.3-0.35 min: 30% B,<br>0.35-1.5 min: 70% B,<br>1.5-1.8 min: 90% B,<br>1.8-2.8 min: 90% B,<br>2.8-2.9 min: 5% B,<br>2.9-4.5 min: 5% B,<br>(A: Solvent A; B: Solvent B) |
| **Injection Volume:** | 1 uL |
| Auto-sampler was used in external rinsing mode, with R0 = 95/5 Water / Acetonitrile and R3 = 1 / 1 IPA / Methanol. |  |

**Table-1.4:** MS MRM Transitions observed (for quantitation):

| Compounds | Ion | Precursor Ion (m/z) | Product Ion (m/z) |
| --- | --- | --- | --- |
| RDV (target, drug): | Quantitation ion | 603.2 | 200.0 (@41) |
|  | Quantitation ion | 603.2 | 229.0 (@21) |
| <sup>13</sup>C<sub>6</sub>-RDV (Internal standard for RDV) | Quantitation ion | 609.2 | 206.0 (@25) |
|  | Quantitation ion | 609.2 | 229.0 (@26) |
| GS-441524 (target, metabolite): | Quantitation ion | 292.1 | 163.0 (@25) |
|  | Quantitation ion | 292.1 | 136.0 (@26) |
| <sup>13</sup>C<sub>5</sub>-GS-441524 (Internal standard for metabolite) | Quantitation ion | 297.1 | 204.1 (@25) |
|  | Quantitation ion | 297.1 | 148.0 (@30) |

### 1.7: Calculation of RDV and GS-441524 from the chromatogram

The ratio of RDV and its isotope, ^13^C_6_-ISTD (as an internal standard) was calculated. Similarly, the ratio of GS and its internal standard, ^13^C_5_-ISTD was also calculated. From that ratio, the amount of RDV and GS were determined using the linear equation derived from their respective standard curve.

The values were normalized using the Dilution factor (2 × 20 × 4=160) used for Original Plasma sample.

## Results and Discussion

### 1: Standard Curve of RDV and GS-441524 (Table-2, Fig. 2)

A representative standard curve for RDV and GS-441524, both were shown in Fig. 2 with R^2^ value (>0.99). The experiment was repeated 3 times with similar results.

### 2: Stability of different RDV test materials in presence of Rat Plasma *in vitro* were shown in Figs 3–5

**Fig. 3:**
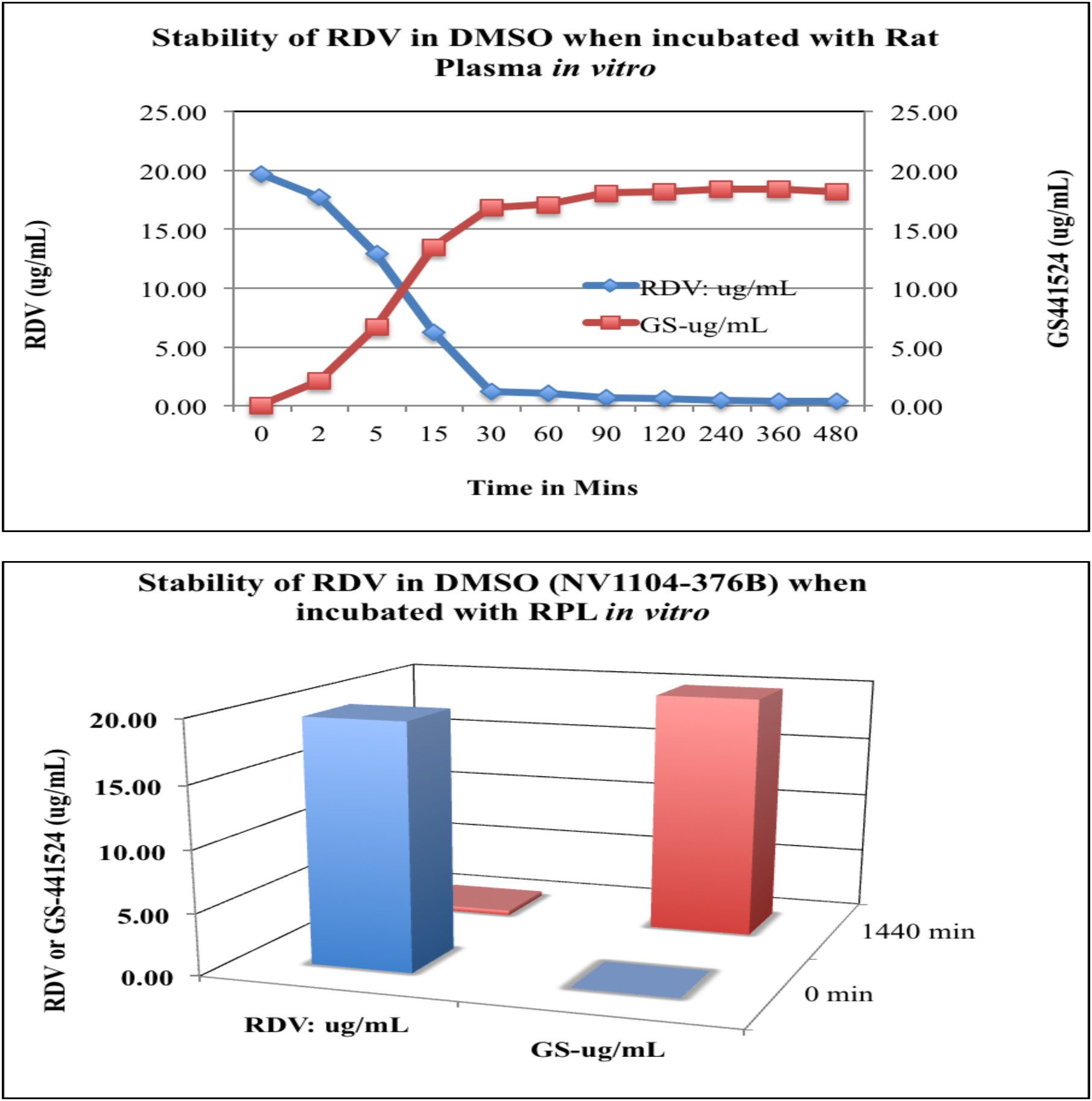
Stability of RDV in DMSO (NV1104-376B) incubated with RPL in vitro: The sample, RDV (in DMSO), was tested for their stability in presence of Rat Plasma in vitro. Methodologies of incubation in reaction mixture, extractions, are same as we did for standard curve determinations of RDV/GS-441524. The results show that RDV alone has a very short life in presence of Plasma. GS-metabolite formations are representative of RDV breakdown, and supportive to each other data. Each data points are the Mean ± SD of three experiments done in duplicate.

**Fig. 4:**
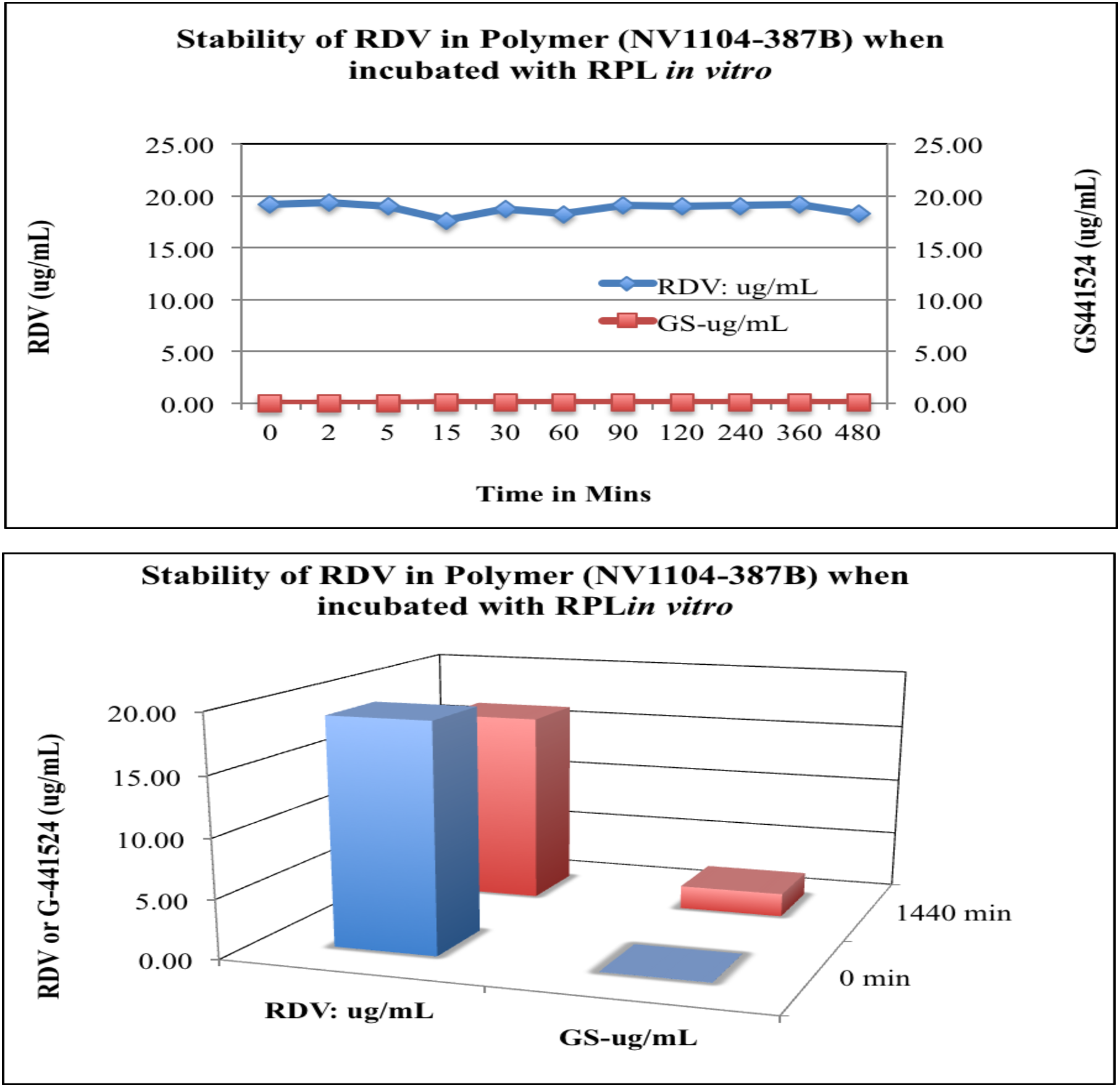
Stability of RDV encapsulated in polymer (NV1104-387B) and incubated with RPL *in vitro:* The sample was tested for their stability in presence of Rat Plasma in vitro. Unlike of RDV in DMSO, we noticed here the protective capacity of polymer encapsulation of RDV from it’s plasma-mediated breakdown. Each data points are the Mean ± SD of three experiments done in duplicate.

**Fig. 5:**
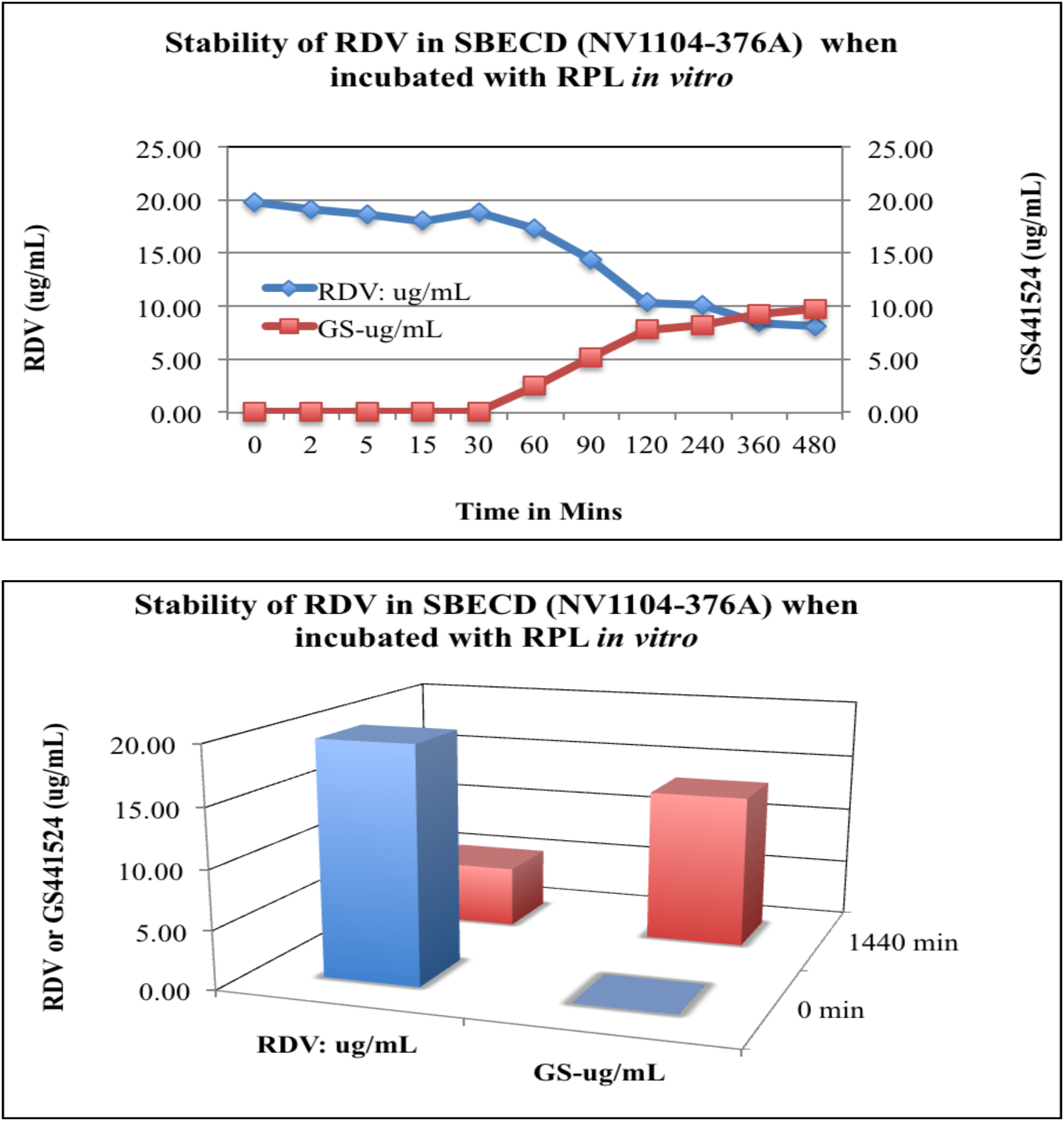
Stability of RDV in SBECD (NV1104-376A; Gilead) in RPL, in vitro: The samples RDV in SBECD (Gilead) were tested for their stability in presence of Rat Plasma in vitro as described in Methodologies section. GS-441524, the RDV metabolite formations are representative of RDV breakdown, and supportive to each other data. Each data points are the Mean ± SD of three experiments done in duplicate.

Test materials are:

i. RDV in DMSO: MeOH (1:9)
ii. RDV-NV387 (RDV encapsulated in NV-387 polymer)
iii. RDV-in SBECD (NV1104-376A, Gilead)

Other test materials used in these experiments are the different vehicles used as a negative controls, such as, DMSO:MeOH (1:9), NV-387 polymer itself, and PBS buffer.

In brief, NV-387 Polymer can encapsulate Remdesivir efficiently, as we see RDV-387 show RDV peak in LC-MS chromatography, where as only the polymer NV-387 does not show any RDV peak (data not shown). We have also tried to measure RDV and GS-441524 level in the respective vehicles, like DMSO, Polymer itself NV-387, and PBS, but the results are below the detection level (data Not shown).

### 3. Comparative analysis of RDV breakdown and GS-441524 production by the above test materials in presence of Rat Plasma, *in vitro*. (Table-3, Fig. 6)

**Fig. 6:**
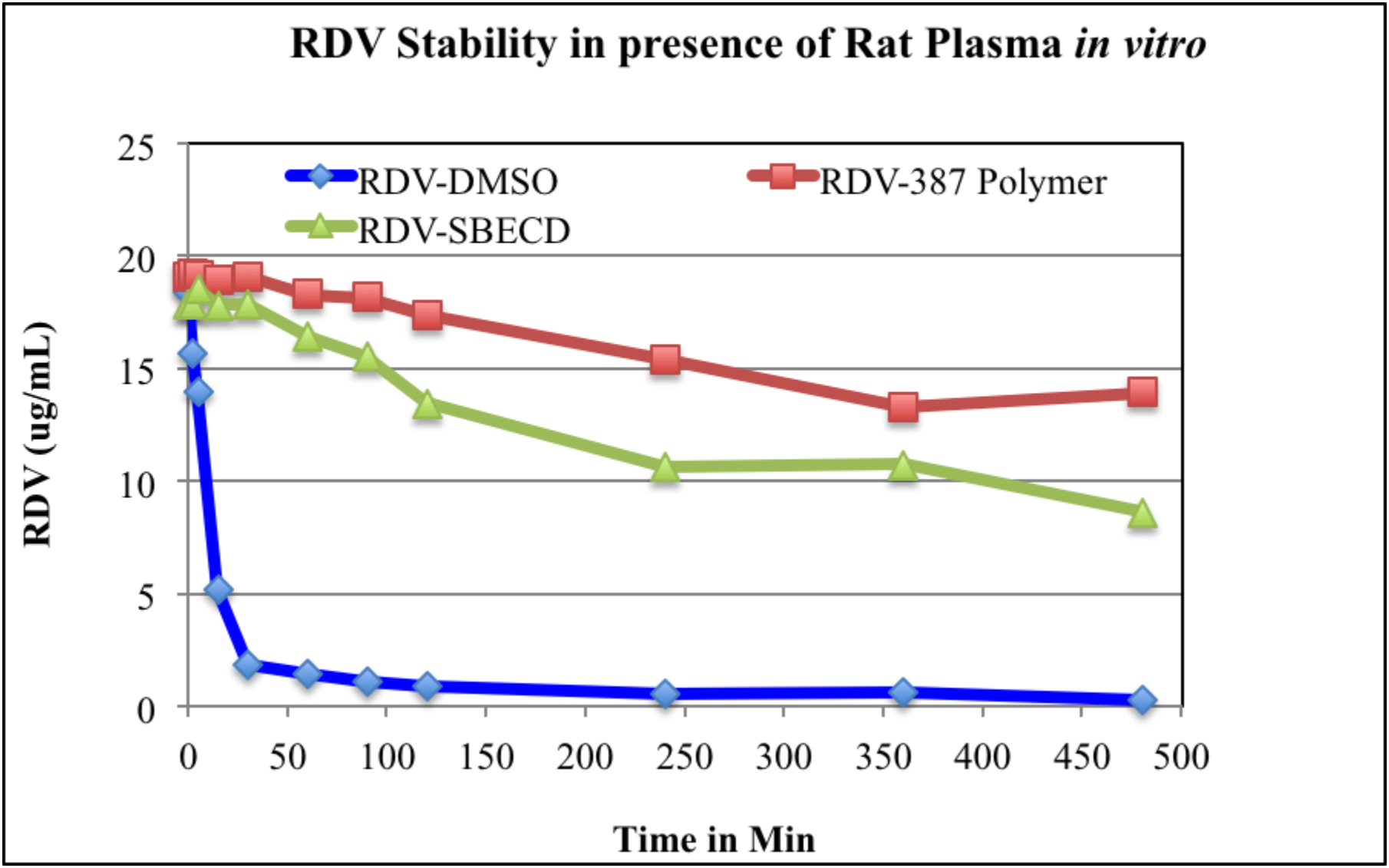
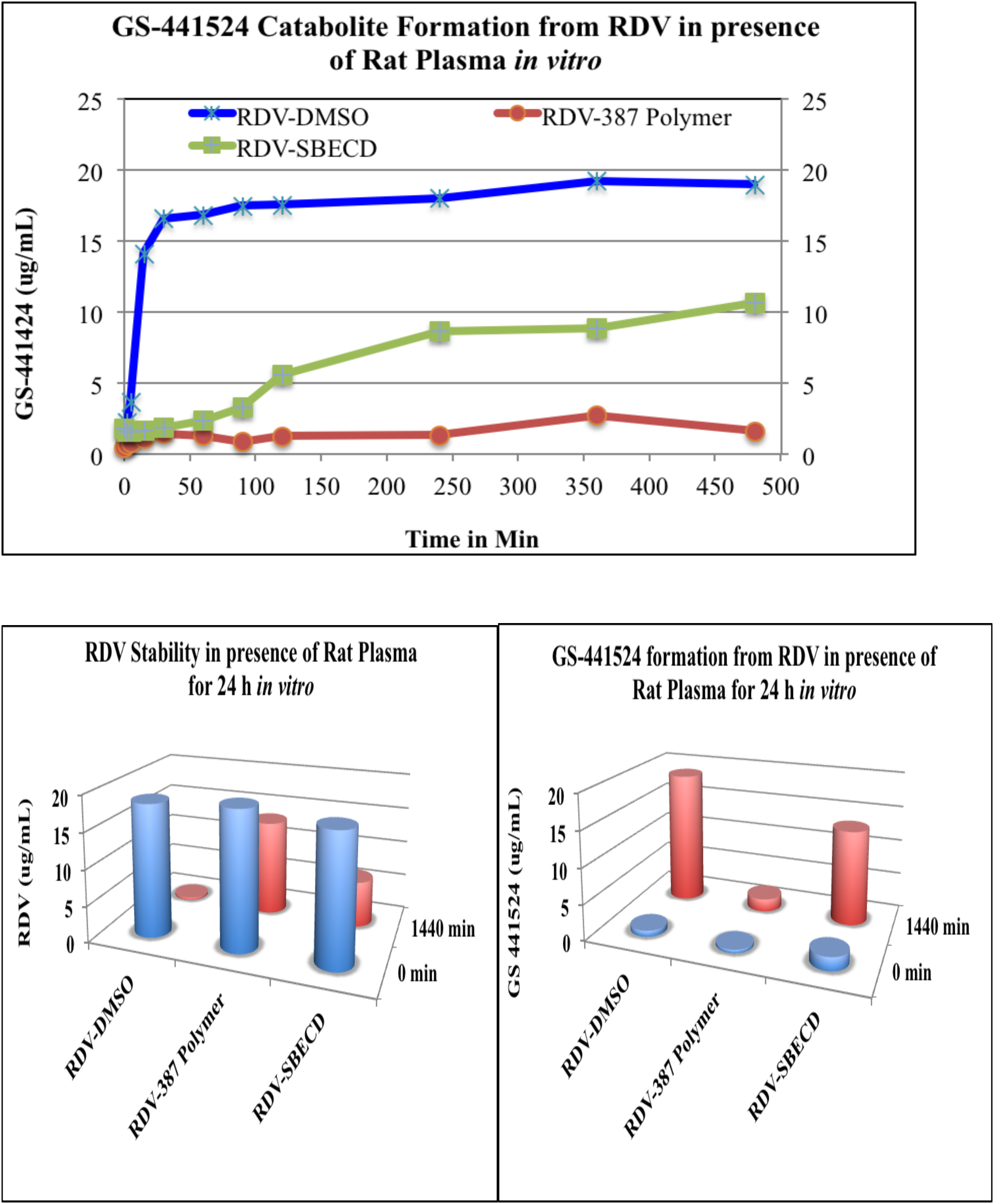
Comparison of Stability of RDV in DMSO, in polymer 387 encapsulated, and in SBECD-In vitro in presence of Rat Plasma: Stability in RPL: The samples, RDV, RDV-encapsulated in MM6 Polymer, RDV in SBECD (Gilead) were tested for their stability in presence of Rat Plasma in vitro. Methodologies of incubation in reaction mixture, extractions, are same as we did for standard curve determinations of RDV/GS. The results show that RDV alone has a very short life in presence of Plasma, but encapsulation in NV-387 polymer make them very stable even after overnight incubation, even compared to Gilead RDV product. GS-441524 metabolite formations are representative of RDV breakdown, and supportive to each other data.

NV-387-polymer encapsulation protects RDV *in vitro* from plasma-mediated catabolism, thus could protect RDV *in vivo* for longer period of anti viral effect even more than that of Gilead-RDV (Figs. 6).

Nonetheless our Platform technology based NV-387-encapsulated-RDV drug has a dual effect on coronavirus, as it being itself posses an antiviral activity **[19, 20],** and also by protecting RDV more than that of Gilead-RDV, renders RDV effectiveness against virus. Further, potential mutations in the virus are unlikely to enable it to escape these drug candidates.

Further, we have already scaled up production of key portions to multi-kilogram scales. We are now initiating an animal model study to finalize two to three best candidates for further testing. We intend to perform certain animal model safety studies, in order to further advance the final candidate for limited human clinical (compassionate) use scenario. This is the fastest timeframe that a drug candidate truly directed at the SARS-CoV-2 has been developed by any Company to date *(https://www.marketwatch.com/press-release/nanoviricides-develops-highly-effective-broad-spectrum-drug-candidates-against-coronaviruses-2020-05-12. Nanoviricides Develops Highly Effective Broad-Spectrum Drug Candidates Against Coronaviruses. Published: May 12, 2020 at 7:15 a.m. ET)*

## Conclusions

It is needless to say that in the present days, rapid design & construction, as well as synthesis and manufacture of anti SARS-CoV-2 has a crying need. Although our approach are very potential for the COVID-19 therapy, but we have been limited to studying effectiveness against available BSL2 level strains of coronaviruses, as they do not cause the severe pathology in humans. No other drug currently available in the development for SARS-CoV-2 virus in the USA, be it antibody, small chemical, or otherwise, has been tested against any coronaviruses at present, to the best of our knowledge.

We are in contact with USAMRIID (United States Army Medical Research Institute of Infectious Diseases) and Robert Davey at NIEDL in Boston, for testing against SARS-CoV-2 itself. We have not received back a Materials Transfer Agreement to enable initiating these studies. Meanwhile its report to scientific community as well as to the common people for their attention and assistance in furthering this endeavor, to contribute to the fight against the SARS-CoV-2 pandemic and position to win the fight.

## Abbreviations

CoV: Coronavirus
HCoV: Human Coronavirus
FDA: Food and Drug Administration
hAPN: human Aminopeptidase N
HAT: Human Airway Trypsin-like protease
SARS: Severe Acute Respiratory Syndrome
IBV: Infectious Bronchitis Virus
TMPRSSII: Transmembrane Protease
MERS: Middle East Respiratory Syndrome
RBD: Receptor Binding Domain
hACE2: human Angiotensin Converting enzyme 2
RNA: Ribonucleic Acid
hDPP4: human Dipeptidyl Peptidase 4

## Authors’ contribution

All the authors contributed equally to prepare this article, read and approved the final manuscript.

## Acknowledgement

We acknowledge all our colleagues, Secretaries for their help during the preparation of the manuscript by providing all the relevant information. Thanks to Ms. Bethany Pond for her editorial assistant. Funds are from Nanoviricide, Inc. company and they are here-by acknowledged.

## Ethical Statement

Not applicable

